# Time-Resolved Phenotyping Reveals Heterogeneous Rice Seed Germination Dynamics in Shallow-Water Culture

**DOI:** 10.64898/2026.08.07.743436

**Authors:** Jinfeng Zhao, Yan Ma

## Abstract

Germination percentage is an endpoint measure and therefore does not describe when an individual seed begins visible growth or how rapidly its radicle and plumule expand. We developed a time-resolved phenotyping workflow to quantify rice seed germination continuously in shallow-water culture. A single industrial camera moved along a 1 m rail and imaged three culture boxes at 1 h intervals for up to 80 h. The archive comprised 1,062 full-frame images and 6,372 seed-level repeated observations under the six-seed field-of-view configuration. A physical grid maintained seed identity through time and enabled individual regions of interest to be extracted. Whole-seed foregrounds were obtained with a pretrained U^2^-Net, and a masked RGB intensity rule separated newly emerging tissue from the darker hull. For each tracked seed, projected emerging-tissue area and interval growth rate were calculated. Three representative normally germinating seeds first showed measurable tissue at 48 h, yet subsequently followed distinct trajectories: final projected areas ranged from 2,605 to 4,700 pixels and peak interval growth rates ranged from 106.88 to 287.92 pixels h^−1^. B-1 accumulated 63.71% of its final visible area during 72–80 h, whereas B-3 accumulated 73.51% during 60–72 h. Thus, seeds with the same observed emergence interval can differ substantially in the timing and magnitude of post-emergence expansion. The workflow converts repeated images into biologically interpretable temporal phenotypes and provides a basis for nondestructive studies of rice seed vigor and germination heterogeneity.

## 1 Introduction

Germination marks the resumption of embryo growth and culminates visibly in radicle protrusion through the surrounding seed tissues [1]. The timing, uniformity, and subsequent expansion of emerging tissues are important components of seed vigor, but these dynamic properties are only partially represented by a single final germination percentage [2]. Conventional testing therefore requires repeated inspection by trained personnel and still yields observations at only a limited number of time points.

Imaging provides a nondestructive record from which the same seed can be measured repeatedly. Early machine-vision systems automated radicle-emergence counting under controlled conditions [3]; later platforms introduced high-throughput curve fitting [4], microplate-based acquisition [5], printable well arrays [6], and deep-learning detection for rice germination-rate assessment [7]. Other image-based systems quantify seed morphology [8], imbibition kinetics [9], or root development and hourly growth [10]. Collectively, these studies show that automated phenotyping can move seed testing from sparse manual counts toward dense temporal measurements.

The present problem is more difficult than endpoint counting. Rice seeds are imaged in shallow water, where the meniscus and the white culture grid generate bright reflections. Daytime natural light and nighttime LED illumination change the intensity and color distribution. The emerging radicle and plumule are narrow, pale structures that may occupy only a few pixels and remain attached to a textured hull. In addition, several seeds appear in each full-frame image and must retain their identities throughout the sequence.

Deep convolutional models have substantially improved image-based plant phenotyping [11]. Reviews of high-throughput plant analysis emphasize that machine learning is most informative when its outputs correspond to interpretable plant traits and are supported by an appropriate experimental design [12, 13]. In this study, image segmentation is therefore used as a measurement step rather than as the biological endpoint. The measured traits are the time of first visible emerging tissue, projected tissue area, interval expansion rate, peak-rate timing, and post-emergence accumulation.

We asked three biological questions. First, can hourly nondestructive imaging delimit the onset of visible germination for an individual rice seed? Second, do seeds with similar emergence timing subsequently differ in the magnitude and timing of tissue expansion? Third, can projected area and interval growth rate provide complementary descriptions of these trajectories? To address these questions, we combined a grid-organized shallow-water culture system with seed-level image analysis and evaluated the recorded 0–80 h trajectories.

## 2 Biological and Imaging Background

### 2.1 Time-Resolved Seed Phenotyping

Ducournau *et al.* used color-image segmentation and automated acquisition to monitor sunflower radicle emergence [3]. Germinator combined experimental design, automatic scoring based on radicle-to-seed-coat color contrast, curve fitting, and parameter extraction [4]. ScreenSeed further coupled hourly imaging of 96-well plates with database-backed analysis for large Arabidopsis screens [5]. Chai *et al.* used a 3D-printed planter and well array to standardize seed placement [6]. These systems illustrate a recurring principle: physical organization of the scene simplifies downstream image analysis.

For cereal crops, SmartGrain demonstrated high-throughput measurement of rice-seed shape and size from digital images [8]. A scanner-based maize platform measured changes in kernel projected area every 10 min and related imbibition parameters to germination [9]. Zhao *et al.* applied deep learning to automatic rice-germination-rate evaluation in dense scenes [7]. MultipleXLab integrated motorized imaging and deep segmentation to monitor seed germination and root growth [10]. The present study complements these systems by resolving the appearance and expansion of emerging rice tissues under a shallow-water germination environment.

### 2.2 Image Analysis as a Phenotyping Tool

U-Net’s skip-connected encoder-decoder architecture preserves localization while incorporating context [14]. Residual learning supports substantially deeper feature extractors [15], while DeepLabv3+ combines atrous multiscale context with decoder-based boundary refinement [16]. U^2^-Net uses nested residual U-blocks to aggregate several receptive-field scales within each stage [17]. Its multiscale structure is attractive for a seed foreground that contains both a large hull and thin roots or shoots.

Modern promptable models such as Segment Anything may reduce the cost of constructing reference masks [18], and data augmentation can improve robustness when training data are limited [19]. Here, however, the purpose of the image model is narrower: it defines the whole-seed foreground so that a biologically interpretable color contrast can quantify newly visible radicle and plumule tissue. The physical grid provides persistent identity, and the temporal output is analyzed at the level of each seed.

## 3 Materials and Methods

### 3.1 Study Design and Phenotypic Readouts

The experiment followed rice seeds from the start of imbibition through visible tissue emergence and subsequent elongation. The primary phenotypic readouts were projected emerging-tissue area and its interval rate of change. Secondary descriptors included the observed emergence interval, peak interval rate, peak-rate time, mean post-detection expansion rate, and the ratio between final area and area at first detection. Figure 1 summarizes the acquisition, seed localization, tissue extraction, and time-series analysis steps.

**Figure 1:**
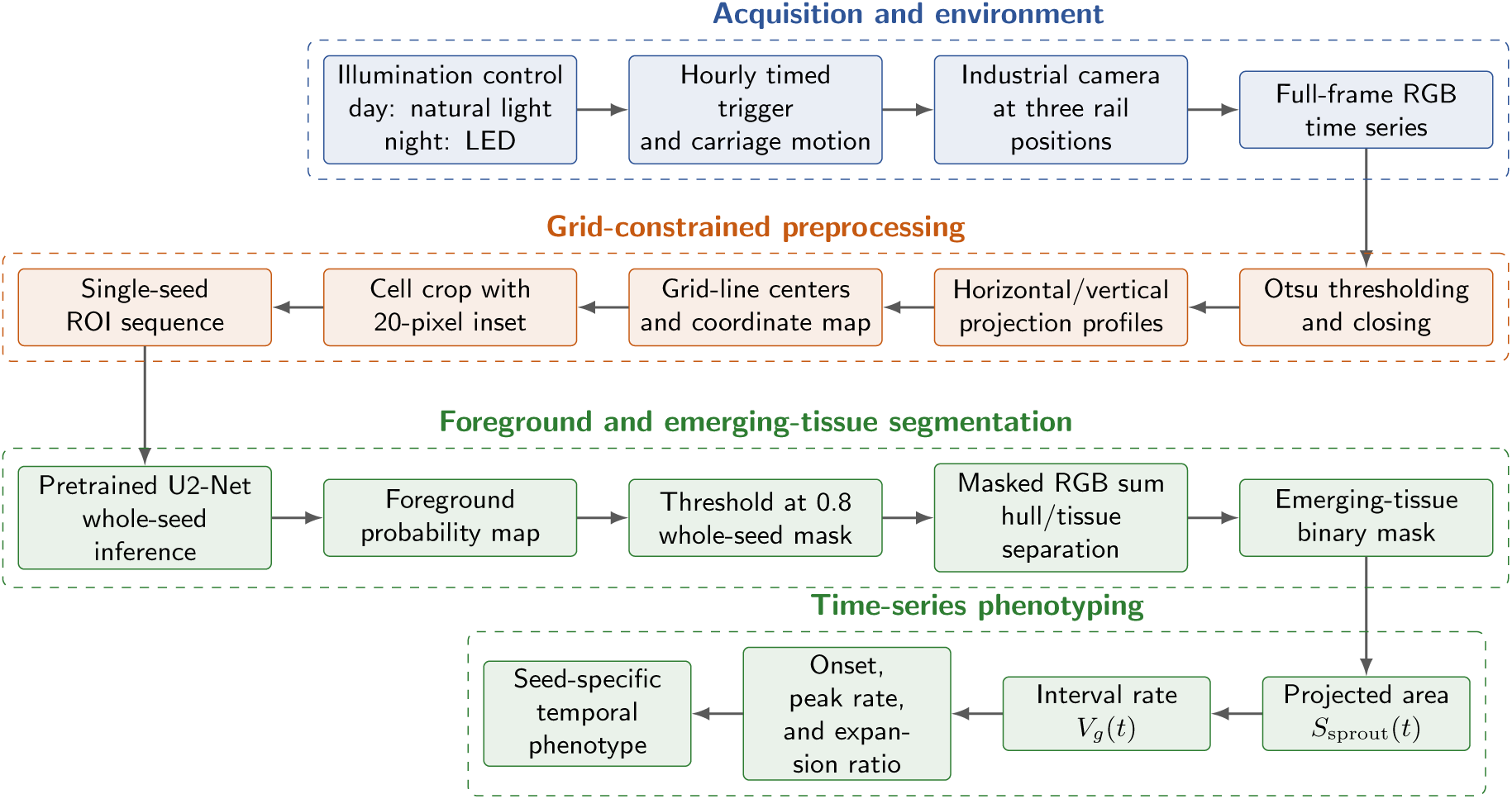
Acquisition-to-phenotype workflow. Physical scene constraints are applied before learned foreground segmentation, and the RGB decision is evaluated only inside the whole-seed mask.

### 3.2 Shallow-Water Culture and Imaging Platform

Rice seeds were placed in three rectangular culture boxes containing a shallow layer of clean water over a dark base. Seeds remained partially submerged during imaging. A white physical grid divided each field of view into repeatable culture cells, maintained the position of each seed, and supplied a stable reference through the sequence. The source record did not retain the rice cultivar or seed-lot identity; the biological observations are therefore reported at the individual-seed level without a cultivar comparison.

The imaging platform consisted of a rigid aluminum frame, a horizontal linear rail with an effective travel distance of 1 m, a motorized carriage, an industrial camera, and a motion controller (Fig. 2). The camera was mounted vertically with its optical axis approximately normal to the culture plane. The carriage moved to predefined positions above the three boxes and triggered acquisition. Use of one camera maintained a common viewing direction and sensor response across the culture positions.

**Figure 2:**
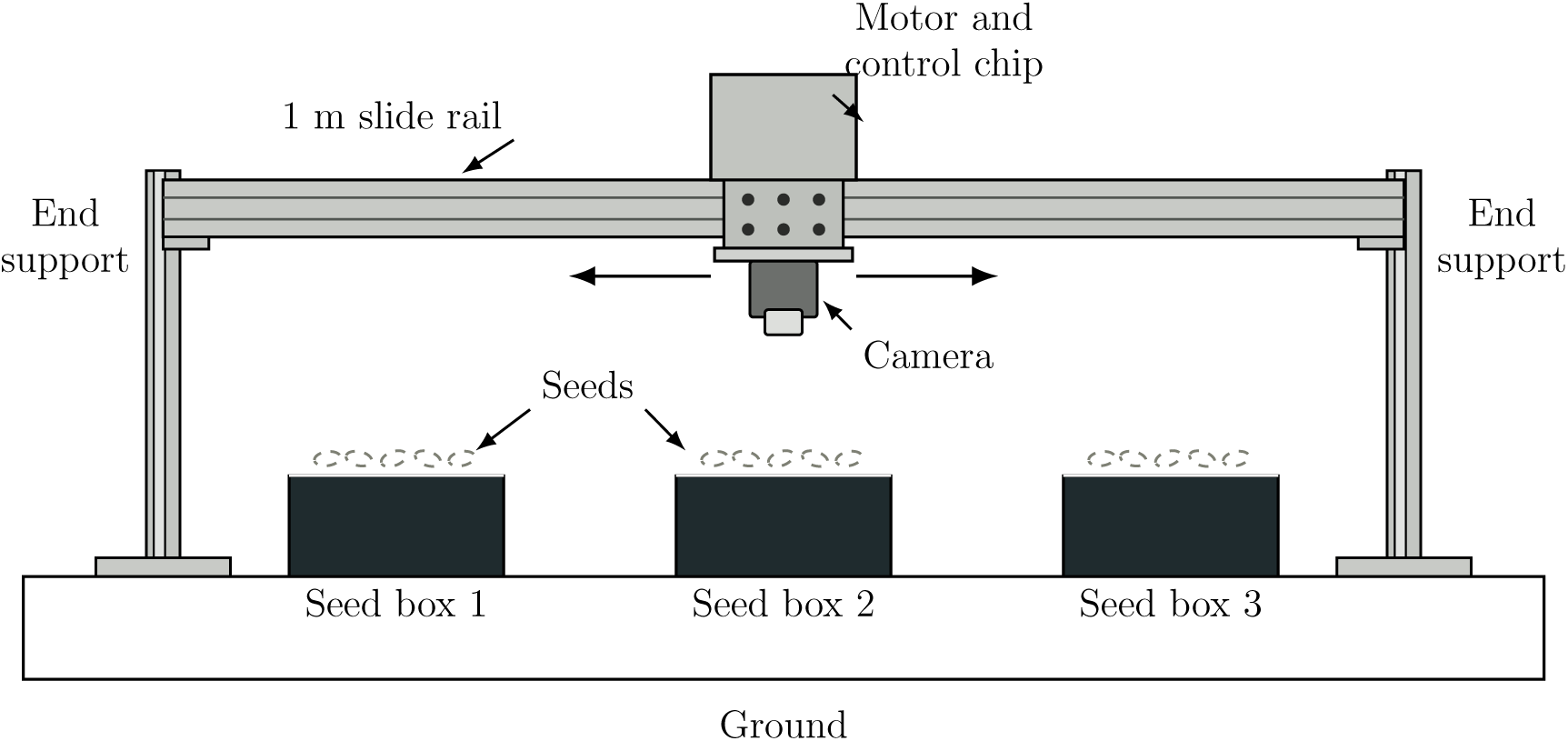
Acquisition platform. A motorized carriage moves the vertically mounted camera along a 1 m slide rail to image three shallow-water seed boxes.

### 3.3 Time-Series Sampling and Observation Set

Images were acquired every 60 min from the start of imbibition until the recorded growth sequence ended, typically at 80 h. From 07:00 to 17:00, the trays were imaged under diffuse natural laboratory light; from 17:00 to 07:00, a constant indoor LED source was used. The mixed-light protocol enabled uninterrupted acquisition but also introduced realistic intensity and color-temperature shifts.

The archive contained 1,062 full-frame RGB images. Under the standard configuration, one full frame contained six seeds, giving

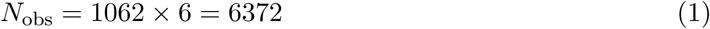

seed-level image observations. These observations are repeated measurements, not 6,372 independent seeds. Analyses of growth dynamics used the three representative normally germinating seeds for which complete selected-time-point measurements were available.

Figure 3 shows representative stages: intact hulls at 0 h, visible pale tissue at 48 h, and elongated radicles and shoots at 80 h. The same white grid remains visible throughout the sequence.

**Figure 3:**
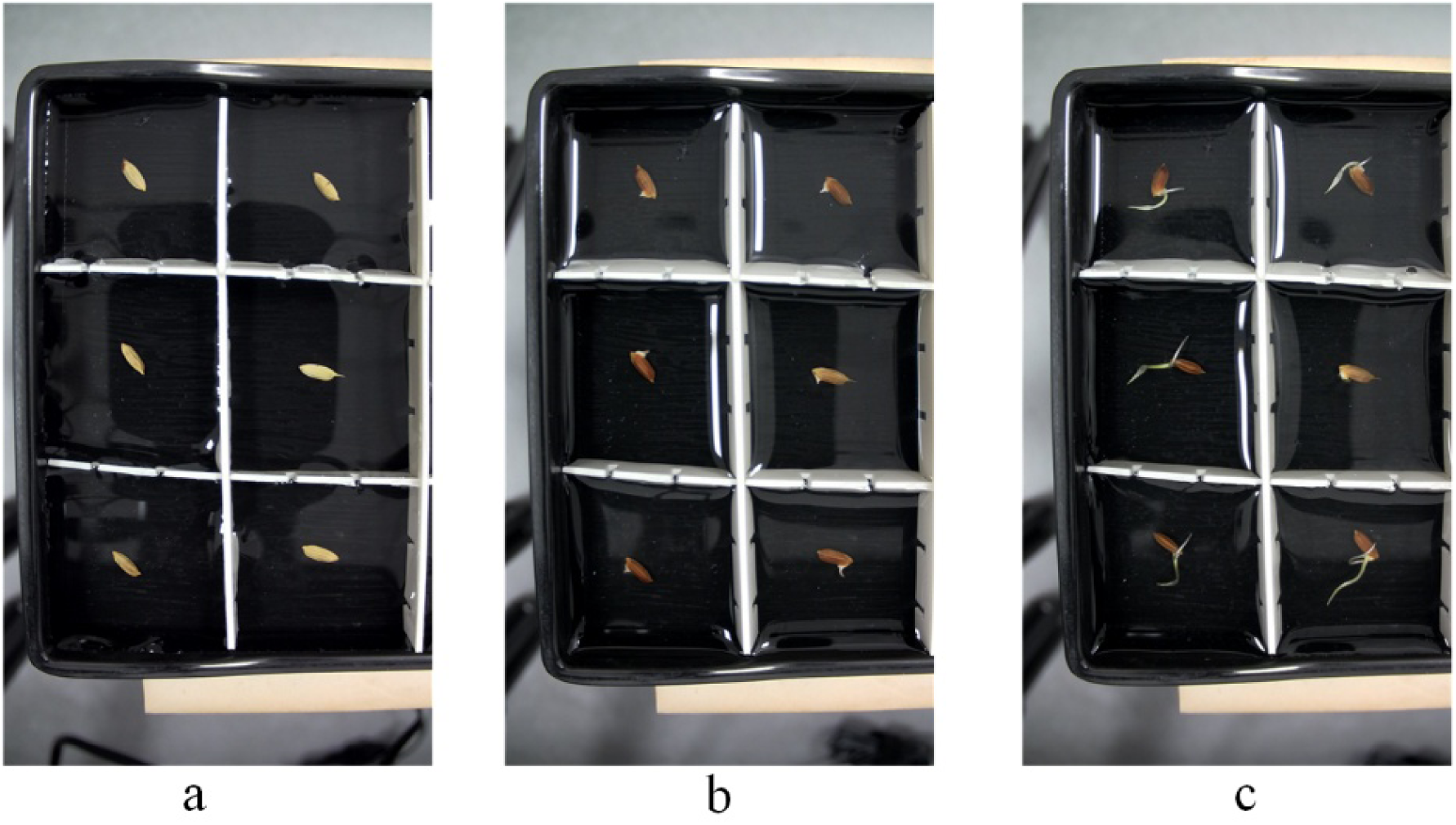
Full-frame images at (a) 0 h, (b) 48 h, and (c) 80 h, showing the transition from imbibition to visible emergence and tissue elongation.

## 4 Image-Based Phenotype Extraction

### 4.1 Grid Localization

Let *I*_RGB_(*x, y*) = [*R*(*x, y*)*, G*(*x, y*)*, B*(*x, y*)] be a full-frame color image and let *I_g_*(*x, y*) be its grayscale conversion. Otsu’s method selects an intensity threshold *T_O_* that maximizes between-class variance [20]. With foreground class probabilities *ω*_0_(*T*) and *ω*_1_(*T*) and class means *µ*_0_(*T*) and *µ*_1_(*T*), the selected threshold is

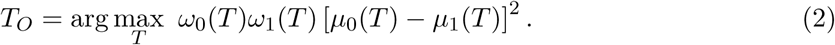

The binary grid candidate is

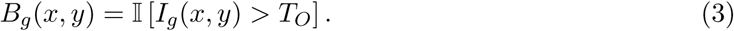

Because the grid is brighter than the dark water background, it dominates *B_g_*. Morphological closing with structuring element *K* connects small discontinuities:

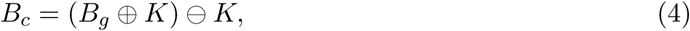

where ⊕ and ⊖ denote dilation and erosion.

Horizontal and vertical projection profiles are computed as

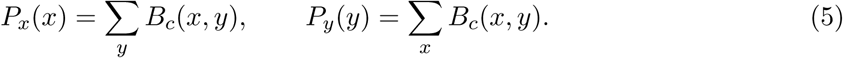

Peak groups in *P_x_*and *P_y_*identify grid-line centers. Adjacent horizontal and vertical centers define rectangular cells. In a production implementation, the detected line set should be checked for the expected number, ordering, minimum spacing, and agreement with the preceding frame before any ROI is accepted.

### 4.2 Persistent ROIs and Reflection Rejection

Once the grid coordinates are established, the same ordered cell map is propagated through the time sequence. For a cell with bounds (*x_l_, x_r_, y_t_, y_b_*), the analyzed ROI is

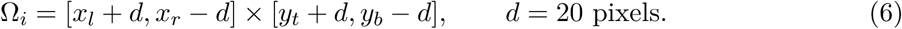

The 20-pixel inset excludes the grid wall and most of the high-intensity meniscus region. It also creates a safety margin against small grid-localization errors. The tradeoff is that a sufficiently long root may eventually cross the inset boundary; such a case should be flagged as truncation rather than silently interpreted as slower growth.

Representative ROIs are shown in Fig. 4. The fixed cell order supplies an identity key (*r, c*) for each seed, so temporal association does not require an additional tracker as long as a seed remains in its assigned cell.

**Figure 4:**
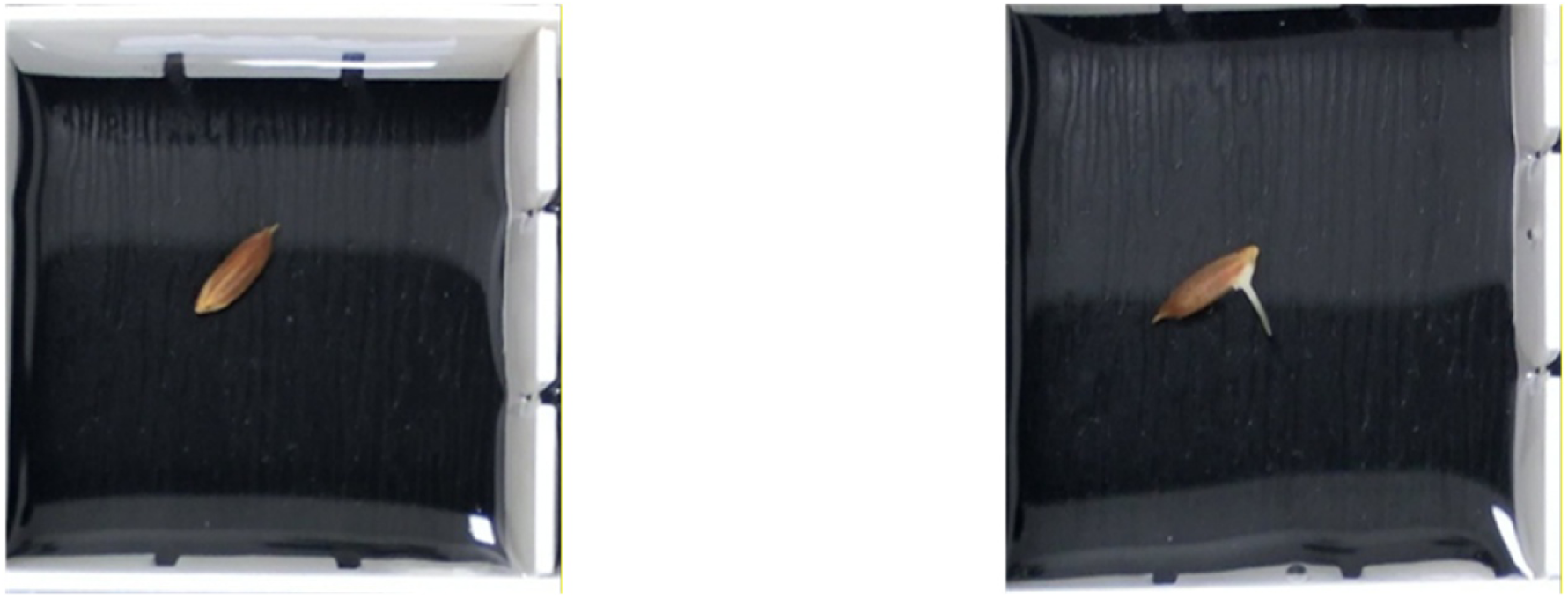
Single-seed ROIs after grid localization and the 20-pixel inward crop. The panels correspond to progressively later stages of germination.

### 4.3 U^2^-Net Whole-Seed Segmentation

The cropped ROI can still contain water texture, nonuniform illumination, and tray-bottom reflections. A pretrained U^2^-Net therefore estimates the complete foreground, operationally defined as the hull plus all connected visible emerging tissues. The network’s residual U-blocks combine local boundary detail with wider spatial context [17]. Residual feature learning is also a standard mechanism for stabilizing deeper vision models [15].

For ROI Ω*_i_* at time *t*, the model returns *p_i,t_*(*x, y*) ∈ [0, 1]. The whole-seed mask is

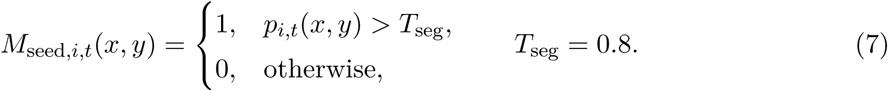

The value 0.8 was used in the experiment. Figure 5 shows representative outputs in which connected elongated structures remain associated with the seed foreground.

**Figure 5:**
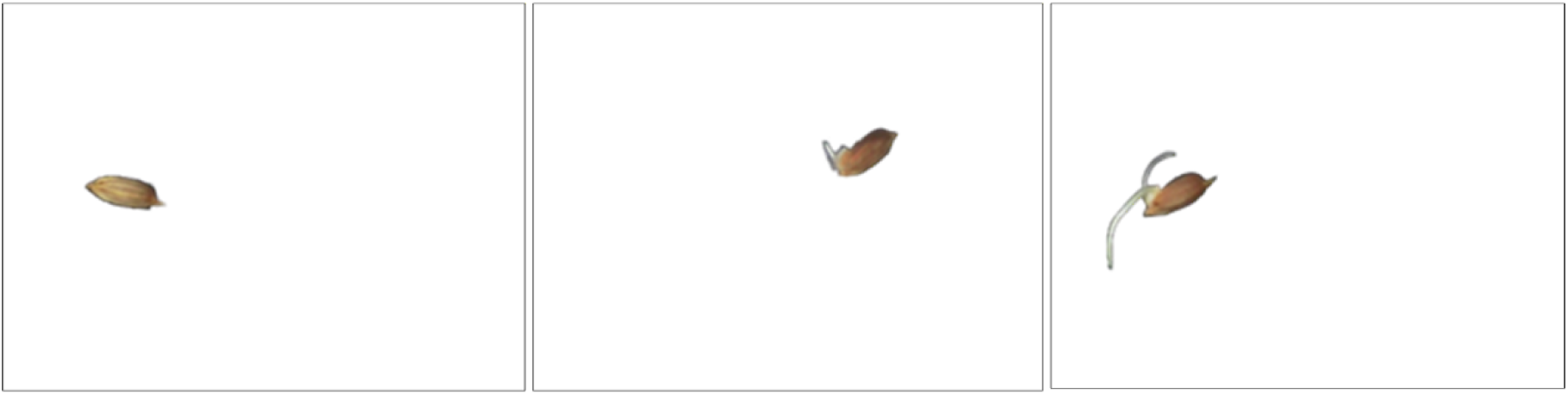
U^2^-Net whole-seed foreground outputs. The masks retain the hull and visible emerging tissues as a single foreground object.

### 4.4 Masked RGB Separation of Emerging Tissue

Within the whole-seed foreground, the hull is generally brown or yellow and the new radicle or plumule is white, pale yellow, or light green. Let

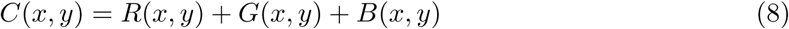

be the RGB channel sum. The emerging-tissue mask is

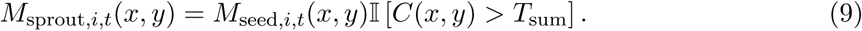

Multiplication by *M*_seed_ prevents bright grid pixels outside the seed foreground from entering the measurement. The experimental record retained the decision rule but not the numerical value or calibration procedure for *T*_sum_.

For reproducible follow-up work, *T*_sum_ should be chosen only on a development set. A practical protocol is to annotate hull and emerging-tissue pixels from several times of day, sweep the threshold over [0, 765], select the operating point that maximizes development-set Dice score or balanced accuracy, and then freeze the threshold before evaluation. DeepLabv3+ [16] and a fine-tuned U^2^-Net should be included as learned baselines. Promptable segmentation [18] may accelerate annotation, but every proposed mask still requires human correction.

### 4.5 Temporal Phenotypes

The projected emerging-tissue area is the foreground-pixel count

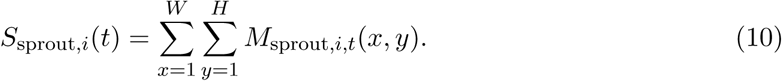

It is reported in pixels because a physical calibration target was not retained. The interval growth rate is the backward finite difference

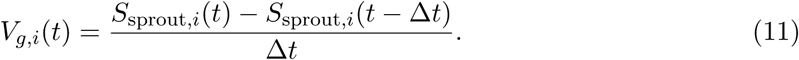

For the complete acquisition sequence, Δ*t* = 1 h. In Table 2, which reports selected time points, Δ*t* is the elapsed time between the displayed observations: 12 h through 72 h and 8 h from 72 to 80 h.

**Table 1:** Summary of Acquisition and Processing Parameters.

| Item | Reported setting | Experimental role |
| --- | --- | --- |
| Imaging platform | One vertically mounted industrial camera on a 1 m horizontal slide rail | Sequential overhead imaging of three seed boxes with a common sensor and viewing direction |
| Culture arrangement | Three rectangular boxes; shallow clean water; dark base; seeds partially submerged | Maintains imbibition while increasing contrast between the rice seeds and the background |
| Spatial organization | White physical grid; six seeds in the standard full-frame field of view | Preserves seed position and enables deterministic single-seed ROI extraction |
| Temporal sampling | One image every 60 min, from 0 h to approximately 80 h | Records the imbibition, visible-emergence, and tissue-elongation stages |
| Daytime illumination | Diffuse natural laboratory light, 07:00–17:00 | Provides near-natural acquisition with realistic intensity variation |
| Nighttime illumination | Constant indoor LED light, 17:00–07:00 | Enables uninterrupted imaging when natural light is unavailable |
| Archived image volume | 1,062 full-frame RGB images and 6,372 seed-level repeated observations | Supplies the time-series image archive processed in this study |
| ROI boundary suppression | 20-pixel inward crop on every side of a detected grid cell | Removes high-intensity meniscus reflections adjacent to the white grid |
| Whole-seed decision | U<sup>2</sup>-Net foreground-probability threshold of 0.8 | Converts the predicted probability map into the binary whole-seed mask |

**Table 2:** Projected Emerging-Tissue Area and Interval Growth Rate at Selected Time Points.

| Time (h) | Area (pixels) |  |  | Rate (pixels h<sup>-1</sup>) |  |  |
| --- | --- | --- | --- | --- | --- | --- |
|  | B-1 | B-2 | B-3 | B-1 | B-2 | B-3 |
| 0 | 0 | 0 | 0 | 0 | 0 | 0 |
| 12 | 0 | 0 | 0 | 0 | 0 | 0 |
| 24 | 0 | 0 | 0 | 0 | 0 | 0 |
| 36 | 0 | 0 | 0 | 0 | 0 | 0 |
| 48 | 572 | 435 | 427 | 47.67 | 36.25 | 35.58 |
| 60 | 734 | 1435 | 693 | 13.50 | 83.33 | 22.17 |
| 72 | 1085 | 1750 | 4148 | 29.25 | 26.25 | 287.92 |
| 80 | 2990 | 2605 | 4700 | 238.13 | 106.88 | 69.00 |

Four additional descriptive quantities are calculated from the selected measurements:

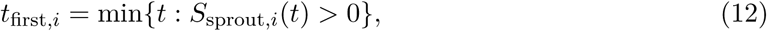

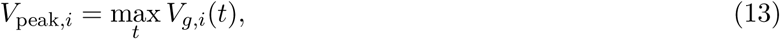

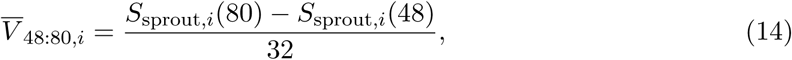

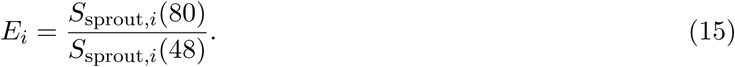

Because the displayed area is zero at 36 h and nonzero at 48 h, visible onset is bounded to (36, 48] h rather than known exactly at 48 h.

### 4.6 Processing Sequence and Quality-Control Flags

For each ordered time point, the processing sequence is:

1. load the RGB frame and localize or verify the white-grid geometry;
2. construct the ordered cell map and apply the 20-pixel inset;
3. infer the U^2^-Net foreground probability for every ROI;
4. threshold at 0.8 to obtain *M*_seed_;
5. evaluate the RGB sum only inside *M*_seed_ to obtain *M*_sprout_;
6. compute area, interval rate, and the seed identity (*r, c*);
7. flag missing grid lines, boundary-touching masks, disconnected high-intensity components, implausibly large area jumps, or a negative area change for manual review.

The deterministic operations are linear in the number of image pixels. Network inference dominates runtime. No runtime benchmark is reported because the archived experiment does not preserve hardware or timing logs.

## 5 Results

### 5.1 Progression from Imbibition to Tissue Elongation

At 0 h, all three illustrated seeds have intact hulls and no visible emerging tissue. At 48 h, pale regions appear near the embryo. By 80 h, roots and shoots extend beyond the hull and produce more complex foreground shapes (Fig. 3). The physical grid remains a stable large-scale structure, and the 20-pixel inset removes most boundary highlights before the neural network operates.

The ROI and segmentation panels in Figs. 4 and 5 show that the processing chain isolates single culture cells and retains the narrow connected structures that appear during germination. This creates a consistent foreground from which visible tissue expansion can be measured through time.

### 5.2 Seeds with Similar Visible Onset Follow Distinct Growth Trajectories

Three representative normally germinating seeds, B-1, B-2, and B-3, were selected for trajectory analysis. Table 2 reports their projected area and interval-rate values. All three remain at zero through 36 h and first become nonzero at the 48 h observation.

The trajectories in Fig. 6 differ in the timing of rapid expansion. B-1 shows modest accumulation through 72 h followed by a late increase to 2,990 pixels. B-2 expands more gradually, with rate increases during both 48–60 h and 72–80 h, and reaches 2,605 pixels. B-3 undergoes the largest mid-to-late pulse, reaching 287.92 pixels h^−1^ at 72 h and 4,700 pixels at 80 h.

**Figure 6:**
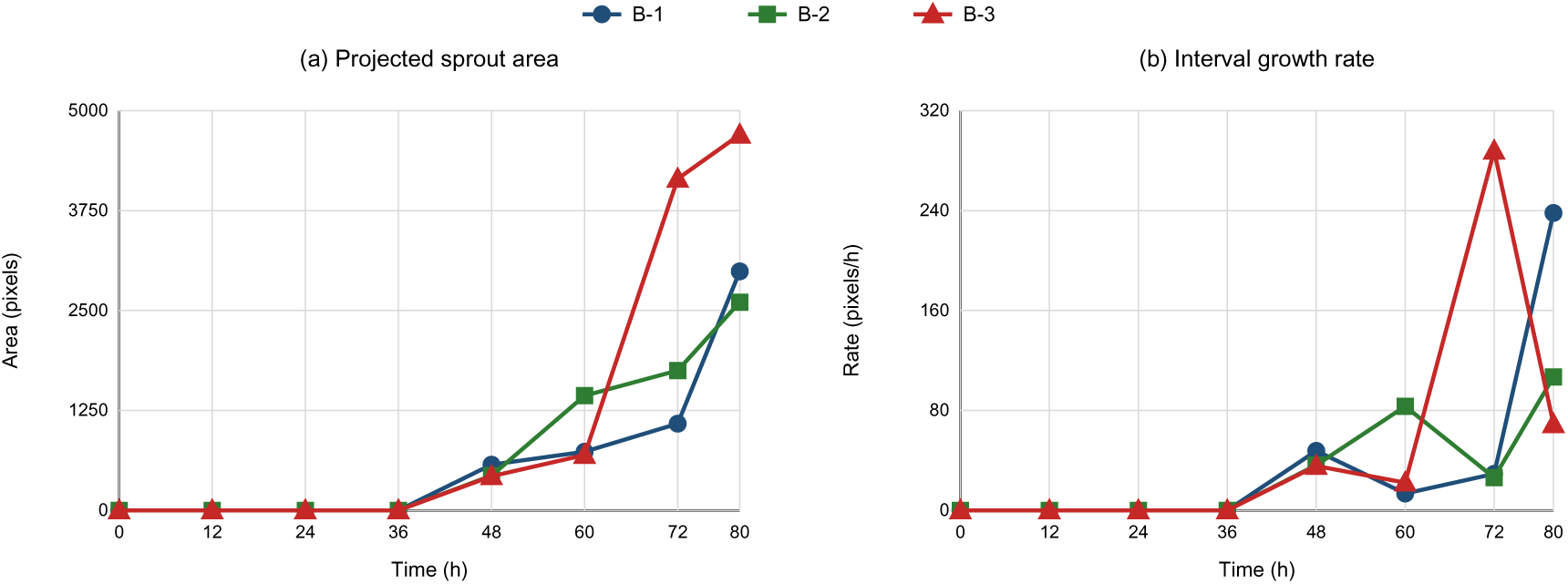
Growth trajectories of three representative rice seeds. Projected area describes cumulative visible expansion, whereas the interval rate emphasizes the timing of growth pulses.

### 5.3 Post-Emergence Expansion Phenotypes

Table 3 adds quantities calculated directly from Table 2. The visible-onset interval is (36, 48] h for every selected seed. B-3 has both the largest post-detection mean expansion rate, 133.53 pixels h^−1^, and the largest expansion ratio, 11.01. B-1 and B-2 peak in the final 72–80 h interval, whereas B-3 peaks earlier, in 60–72 h. Thus, equal first-detection times do not imply equal subsequent vigor trajectories.

**Table 3:** Descriptive Quantities Derived from the Selected Time Points.

| Seed | Visible-onset interval (h) | Final area (pixels) | Peak rate (pixels h<sup>-1</sup>) | Peak time (h) | Post-detection mean rate (pixels h<sup>-1</sup>) / expansion ratio |
| --- | --- | --- | --- | --- | --- |
| B-1 | (36, 48] | 2990 | 238.13 | 80 | 75.56 / 5.23 |
| B-2 | (36, 48] | 2605 | 106.88 | 80 | 67.81 / 5.99 |
| B-3 | (36, 48] | 4700 | 287.92 | 72 | 133.53 / 11.01 |

These descriptors remain sample-specific. The three seeds were selected as representative examples rather than as a randomized biological replicate set, so no hypothesis test or cultivar comparison is justified.

### 5.4 Interval-Resolved Area Accumulation

Table 4 converts every consecutive pair of reported time points into an absolute area gain and its contribution to the final 80 h area. This interval decomposition makes the contrasting growth modes quantitatively explicit. B-1 accumulates 63.71% of its final visible area during 72–80 h. B-2 distributes its largest gain across 48–60 h (38.39%) and 72–80 h (32.82%). In contrast, B-3 accumulates 73.51% of its final area during 60–72 h, consistent with its earlier rate peak.

**Table 4:** Interval-Resolved Emerging-Tissue Area Gains.

| Interval (h) | Area gain (pixels) |  |  | Share of final 80 h area (%) |  |  |
| --- | --- | --- | --- | --- | --- | --- |
|  | B-1 | B-2 | B-3 | B-1 | B-2 | B-3 |
| 0–36 | 0 | 0 | 0 | 0.00 | 0.00 | 0.00 |
| 36–48 | 572 | 435 | 427 | 19.13 | 16.70 | 9.09 |
| 48–60 | 162 | 1000 | 266 | 5.42 | 38.39 | 5.66 |
| 60–72 | 351 | 315 | 3455 | 11.74 | 12.09 | 73.51 |
| 72–80 | 1905 | 855 | 552 | 63.71 | 32.82 | 11.74 |

### 5.5 Projected Area as a Germination Phenotype

The reported area is a two-dimensional phenotype of visible emerging tissue. It combines the radicle and plumule when both are visible and does not represent dry mass or three-dimensional length. Orientation changes can alter projected area even when biological length continues to increase. Its principal advantage is that it can be measured nondestructively from the same seed throughout germination. The cumulative area curve describes visible tissue accumulation, whereas the interval derivative emphasizes transitions between slower and faster expansion.

## 6 Discussion

### 6.1 Temporal Phenotypes Extend Endpoint Germination Scoring

An endpoint germination percentage records whether a seed has crossed a predefined biological criterion. The time-resolved traits used here add two other dimensions: when emerging tissue first becomes visible and how that tissue expands afterward. The three selected seeds shared the same observed onset interval, (36, 48] h, but differed strongly in their final area, peak rate, and peak timing. Emergence time alone therefore did not capture the full phenotypic variation present in these trajectories.

### 6.2 Distinct Modes of Post-Emergence Expansion

B-1 displayed a late-expansion pattern, accumulating almost two-thirds of its final visible area during 72–80 h. B-3 displayed an earlier pulse, accumulating nearly three-quarters of its final area during 60–72 h before its interval rate declined. B-2 showed a more distributed pattern across the post-emergence intervals. These patterns demonstrate that temporal allocation of growth can differ even among seeds that first become visibly positive at the same sampled time point. They provide candidate quantitative traits for comparing seed lots, treatments, or environmental conditions in experiments with biological replication.

### 6.3 Relationship to Existing Seed-Phenotyping Systems

Like Germinator and ScreenSeed [4, 5], this workflow benefits from regular specimen placement and repeated imaging. Like the maize imbibition platform [9], it converts changes in projected area into temporal curves. Its distinguishing feature is the combination of shallow-water culture, persistent grid-defined seed identity, and a foreground model that retains connected emerging tissues. MultipleXLab similarly integrates motorized imaging and deep segmentation for root and germination phenotyping [10]; the present implementation focuses specifically on the onset and expansion of visible rice tissues.

The physical and computational components serve the biological measurement. The grid preserves identity, the inward crop suppresses a repeatable optical artifact, U^2^-Net defines the hull-plus-tissue foreground, and the masked color rule quantifies emerging tissue. This modular design keeps the resulting phenotypes interpretable and allows the same individual to be followed without destructive sampling.

### 6.4 Prospects for Larger Seed-Vigor Studies

The traits reported here can be applied to experiments that explicitly compare cultivars, seed lots, priming treatments, temperature regimes, or water conditions. Such studies can combine the onset interval, area accumulation, peak rate, and peak timing with conventional germination percentage and seedling measurements. Physical image calibration would further allow projected area to be expressed in metric units.

Longer image sequences also create opportunities for models that learn from the trajectory of the same seed. Contrastive temporal learning [21] and multi-view fusion of color, shape, area, and mask confidence [22] provide possible computational extensions. Their biological value would be assessed by whether they improve prediction of independently measured seed-vigor outcomes.

## 7 Conclusion

Hourly nondestructive imaging resolved rice seed germination as a temporal process rather than a single endpoint. A grid-organized shallow-water culture system preserved individual identity, and image-based tissue extraction converted each sequence into projected area and interval growth rate. The three representative seeds first showed measurable emerging tissue within the same (36, 48] h interval, yet their final areas and peak expansion times differed markedly. Interval analysis distinguished a late-expansion pattern in B-1, a distributed pattern in B-2, and an earlier rapid-expansion pattern in B-3. These results show that time of visible emergence and subsequent tissue expansion are complementary phenotypes and support the use of longitudinal imaging for studies of rice seed vigor and germination heterogeneity.

## Data and Code Availability

The image data and analysis code supporting this study are available from the corresponding author upon reasonable request.

## Ethics Statement

This study involved plant seeds only and did not involve human participants or vertebrate animals.

